# C-terminal determinants for RNA binding motif 7 protein stability and RNA recognition

**DOI:** 10.1101/2022.05.13.491737

**Authors:** Amr M. Sobeh, Catherine D. Eichhorn

## Abstract

The 7SK ribonucleoprotein (RNP) is a critical regulator of eukaryotic transcription. Recently, RNA binding motif 7 (RBM7), which contains an RNA recognition motif (RRM), was reported to associate with 7SK RNA and core 7SK RNP protein components in response to DNA damage. However, little is known about the mode of RBM7-7SK RNA recognition. Here, we recombinantly expressed and purified RBM7 RRM constructs and found that constructs containing extended C-termini have increased solubility and stability compared to shorter constructs. To identify potential RBM7-7SK RNA binding sites, we analyzed deposited data from *in cellulo* crosslinking experiments and found that RBM7 crosslinks specifically to the distal region of 7SK stem-loop 3 (SL3). Electrophoretic mobility shift assays and NMR chemical shift perturbation experiments showed weak binding to 7SK SL3 constructs *in vitro*. Together, these results provide new insights into RBM7 RRM folding and recognition of 7SK RNA.

**Graphical abstract:** 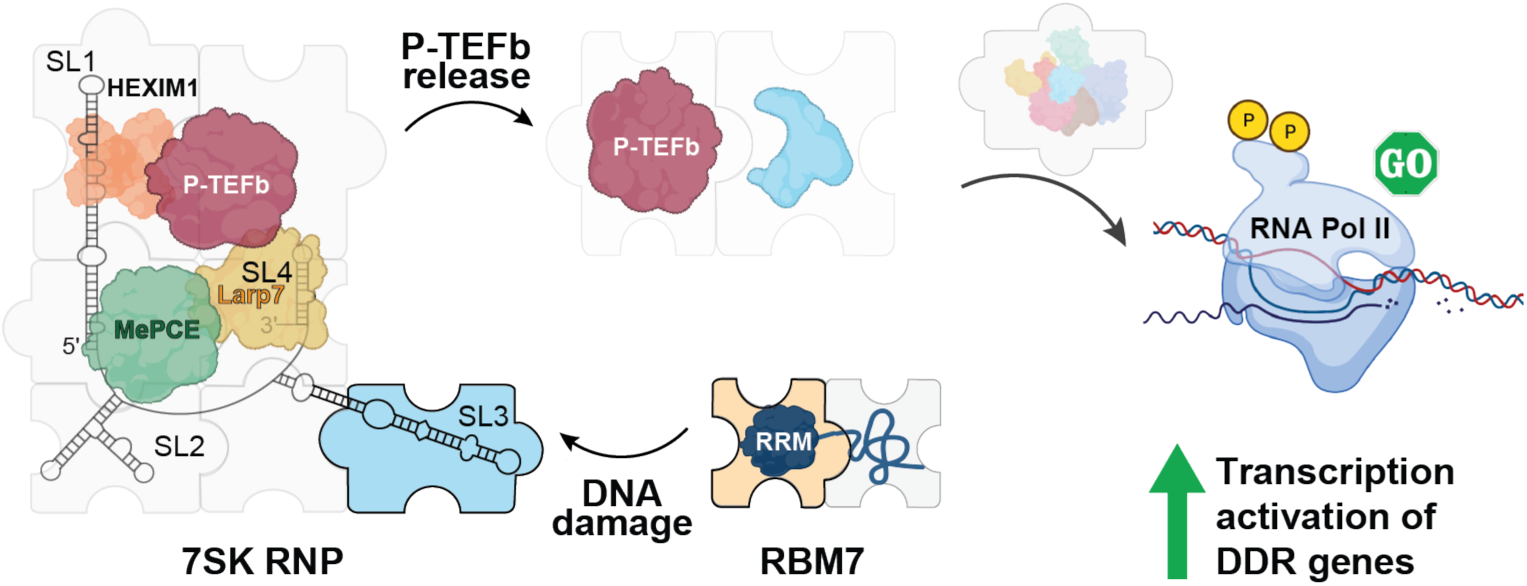

**Highlights:**

- Extending the RBM7 RRM C-terminus improves protein expression and solubility
- The RBM7 RRM interacts weakly with 7SK stem-loop 3 single-stranded RNA
- Solution state NMR dynamics measurements support presence of additional β4’ strand

## Introduction

Proper regulation of RNA polymerase II (Pol II) transcription is required to maintain cellular homeostasis and efficient gene expression [1]. Promoter-proximal pausing of Pol II ∼20–60 nt downstream of the transcription start site is an essential checkpoint in transcription regulation [2-5], controlled in part by the kinase positive transcription elongation factor b (P-TEFb) [5-8]. In metazoa, the 7SK ribonucleoprotein (RNP) sequesters P-TEFb in a catalytically inactive pool and P-TEFb must be released from 7SK RNP to restore P-TEFb kinase activity [9-11].

The 7SK RNP assembles into a core complex comprised of the 7SK RNA, a 331 nucleotide (nt) noncoding RNA in humans [12]; the methylphosphate capping enzyme (MePCE), which caps and remains bound to the 7SK RNA 5’ end [13, 14]; and the La-related protein 7 (Larp7), which binds to the 7SK RNA 3’ end [15-18]. MePCE acts cooperatively with Larp7 to assemble with 7SK RNA to form a stable ternary complex [13, 19]. The accessory hexamethylene bisacetamide-inducible protein 1 or 2 (HEXIM1/2) is required to assemble P-TEFb onto the 7SK RNP and inhibit P-TEFb kinase activity [20-22]. Together, these components minimally constitute the P-TEFb-inactivated 7SK RNP [16, 23, 24]. Numerous accessory proteins have been reported to promote release of P-TEFb from 7SK RNP [5, 25-34]. On release P-TEFb, Bromodomain 4 (Brd4), and other transcription factors trigger the transition of Pol II from a promoter-proximal paused to a processive elongation state [5, 35, 36]. P-TEFb dysregulation or 7SK RNP malfunction is associated with several diseases including blood and solid tumor cancers, cardiac hypertrophy, and primordial dwarfism [5, 37]. Moreover, several viruses manipulate host 7SK RNP for viral survival, notably HIV-1 [38] and more recently SARS-CoV-2 [39].

Spatial organization of 7SK RNP on chromatin may enable a rapid cellular response to environmental conditions. In support of this model, genome-wide analyses report the presence of 7SK RNP at the promoters of protein-coding genes [33, 40, 41]. UV exposure results in the global release of promoter-proximally paused Pol II [42], stimulates the rapid release of P-TEFb from 7SK RNP, and induces activation of UV-response genes [2, 34, 36]. 7SK RNA knockout HAP1 cells showed reduced viability after UV irradiation, with impaired activation of UV-response genes requiring 7SK RNP−P-TEFb to rescue activation [43]. In HEK293 and HeLa cells, exposure to a chemical mimetic of UV-induced DNA damage (4-nitroquinoline 1-oxide, 4-NQO) similarly resulted in decreased viability [44, 45]. 4-NQO exposure induces phosphorylation of the RNA binding protein 7 (RBM7) by the p38^MAPK^−MK2 pathway [34, 46] and transcription activation of genes involved in DNA damage response (DDR) [34, 47, 48]. RBM7 individual-nucleotide resolution UV crosslinking and immunoprecipitation (iCLIP) experiments found that RBM7 interacts with 7SK RNA, and co-immunoprecipitation experiments found that RBM7 interacts with core 7SK RNP proteins MePCE and Larp7 as well [34]. A model has been proposed in which genotoxic stress stimulates phosphorylation of RBM7 to promote release of P-TEFb from 7SK RNP [34]. More recently, p38^MAPK^−MK2-induced phosphorylation of RBM7 has been shown *in cellulo* after SARS-CoV-2 viral infection [39, 49], suggesting a similar pathway may be activated during immune response.

The primary identified function of RBM7 is as a component of the nuclear exosome targeting complex (NEXT), a trimeric complex containing RBM7, the zinc-knuckle protein ZCCHC8, and the RNA helicase hMTR4 [50, 51]. NEXT interacts robustly with the RNA cap-binding complex to induce transcription termination; in addition, NEXT is involved in the degradation of aberrant Pol II transcripts [52-54]. The role of RBM7 in 7SK RNP-mediated DDR is an alternate regulatory mechanism independent of function in NEXT. When phosphorylated, RBM7 has globally reduced interaction to RNA with reduced NEXT function [55]. Importantly, 7SK RNA abundance is not reduced after 4-NQO exposure, indicating that RBM7 is functioning independently of its role in NEXT [34].

Together, these data suggest that RBM7 plays an essential role in the release of P-TEFb from the 7SK RNP complex to respond to genotoxic stress [34]. However, little is known regarding how RBM7 assembles with 7SK RNA and core 7SK RNP protein components. RBM7 contains an N-terminal canonical RNA recognition motif (RRM) with conserved RNP1 and RNP2 sequences [56, 57] (**Fig. 1A-B**) and a C-terminal serine-rich domain that is phosphorylated by p38^MAPK^ [34, 46, 56, 57]. *In vitro*, the RBM7 RRM preferentially interacts with polypyrimidine repeat sequences with micromolar binding affinity [56]. Four high-resolution structures have been determined of the RBM7 RRM by solution NMR spectroscopy or X-ray crystallography, with varying N- and C-terminal ends and observed structural features [56, 58, 59]. Here, we recombinantly expressed and purified RBM7 RRM constructs with various N-terminal fusion tags and C-terminal ends. Using solution NMR spectroscopy, we found that constructs containing longer C-terminal ends have increased solubility and stability compared to shorter constructs. ^1^H-^15^N heteronuclear NOE experiments reporting internal dynamics of the RRM domain support the presence of an additional β-strand observed in X-ray crystallographic structures but not the solution NMR structure. To identify potential RBM7-7SK RNA binding sites, we analyzed deposited data from *in cellulo* crosslinking experiments and performed electrophoretic mobility shift assays and NMR chemical shift perturbation experiments. We found that RBM7 crosslinks specifically to the distal region of 7SK stem-loop 3 (SL3) and binds weakly to 7SK SL3 constructs. Together, these data provide new insights into RBM7 RRM folding and recognition of the 7SK RNA.

**Figure 1.**
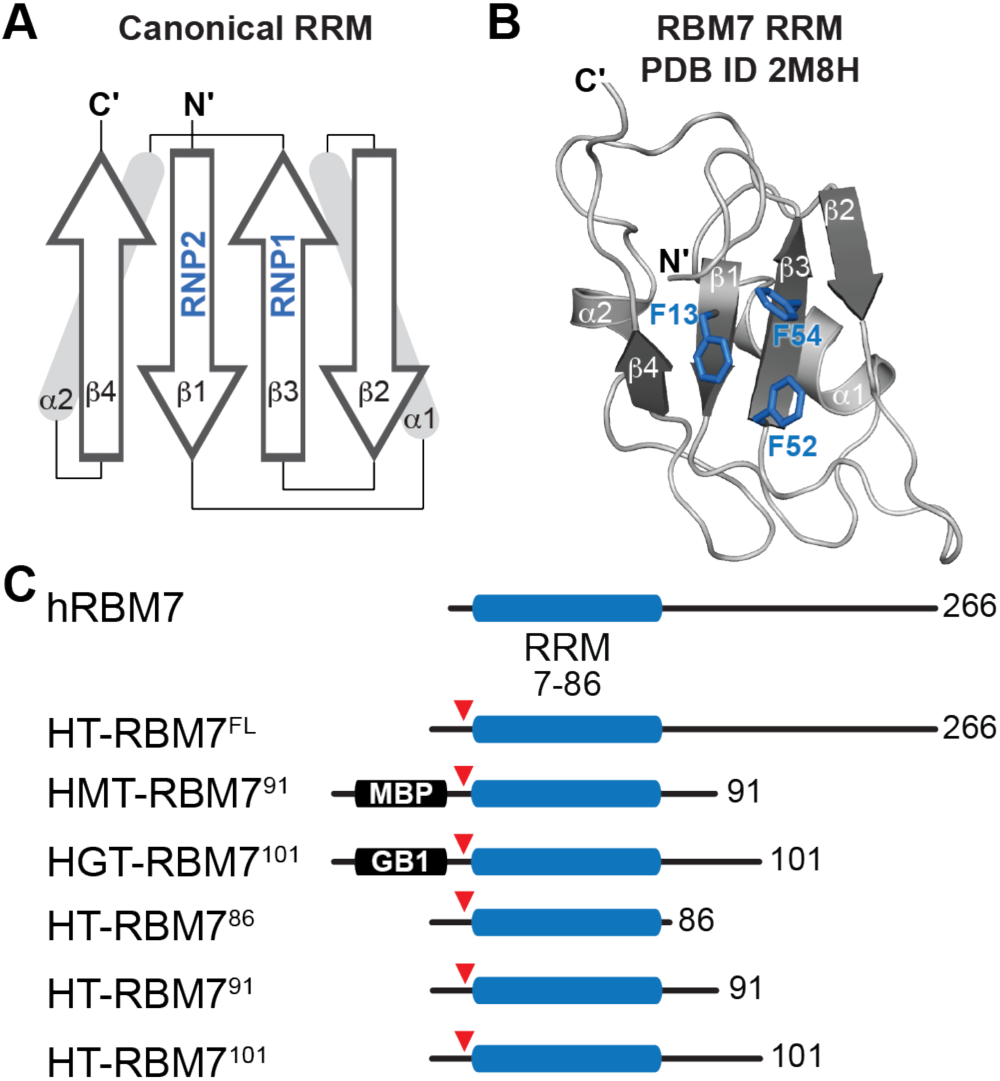
RBM7 domain topology and constructs used in study. A) Schematic of canonical RNA recognition motif (RRM) with conserved RNP1 and RNP2 shown in blue. B) Solution NMR structure of human RBM7 RRM (PDB ID 2M8H) with conserved Phe residues in RNP1 and RNP2 shown as blue sticks. C) Constructs used in study. RRM shown in blue and TEV cleavage site indicated with red triangle.

## Methods

### Recombinant protein construct design, expression, and purification

The DNA sequence encoding human RBM7 was codon optimized for *E. coli* codon bias and purchased as a gblock (Integrated DNA Technologies). The RBM7 gene was cloned into a pET vector containing N-terminal His_6_, tobacco etch virus (TEV) protease recognition site, and maltose binding protein (MBP) domains with a kanamycin resistance gene (Addgene plasmid #29656) using *in vivo* assembly [60]. Briefly, Benchling was used to design overlapping primers complementary to the 5’ and 3’ ends of both the gene insert and vector insertion sites. PCR was performed to produce gene insert and vector with complementary 5’ and 3’ ends. PCR products were purified using a PCR cleanup kit (Zymo Research) and directly transformed into DH5α competent cells. Individual colonies were selected and cultured in LB media with kanamycin, followed by plasmid purification using a plasmid miniprep kit (Qiagen). All constructs derived from this plasmid were generated using site-directed mutagenesis with the exception of the GB1 domain, which was inserted using *in vivo* assembly. Sequences were verified using Eurofins Genomics LLC.

For protein expression, plasmids were transformed into *E. coli* NiCo21(DE3) competent cells (New England Biolabs). Cells were cultured with shaking in M9 minimal media with kanamycin at 37 °C to an OD_600_ of 0.6-0.8, transferred to 18 °C, expression induced with 0.5 mM isopropyl β-D-1-thiogalactopyranoside (IPTG), and grown for 18-20 h. For NMR experiments, ^15^N-labelled ammonium chloride (Cambridge Isotope Laboratory) was used in the M9 media. Buffers used for protein purification were adapted from [56]. Cells were harvested by centrifugation and resuspended in buffer R (50 mM NaPO_4_ pH 7.0, 1 M NaCl, 40 mM imidazole, 10 mM MgCl_2_, 10% glycerol, 1 mM tris(2-carboxyethyl)phosphine (TCEP)), 0.5 mM PMSF, and lysozyme. After sonication, soluble and insoluble fractions were separated by centrifugation at 35,000 xg for 45 min, and the supernatant was clarified by 0.45 µm syringe filter. Affinity purification was performed using nickel nitriloacetic acid (Ni-NTA) affinity column (Qiagen) attached to an ÄKTA start system (GE Healthcare). After loading the supernatant, the column was washed with 10-20 column volumes (CV) of buffer R followed by elution in a linear gradient with buffer E (50 mM NaPO_4_ pH 7.0, 250 mM NaCl, 300 mM imidazole, 10% glycerol, 1 mM TCEP). Fractions were analyzed by 4-12% Bis-Tris NuPAGE™ gels (ThermoScientific). To remove the His_6_-MBP or His_6_-GB1 tag, protein was dialyzed against RBM7 storage buffer (20 mM NaPO_4_ pH 5.5, 50 mM L-glutamate, 50 mM L-arginine and 1 mM TCEP) in the presence of TEV protease (Addgene #92414 [61] recombinantly expressed and purified in house) for 2-3 hours at room temperature. After dialysis, cleaved RBM7 protein was purified from His_6_-MBP or His_6_-GB1 by a second Ni–NTA purification step (Qiagen).

As a final purification step, proteins were purified by size exclusion chromatography using a HiLoad16/600 Superdex 75 pg (GE Healthcare) column attached to a ÄKTA Prime M25 FPLC system (GE Healthcare) in RBM7 storage buffer. Fractions were analyzed by 4-12% Bis-Tris NuPAGE™ gel. Pure fractions were concentrated using a 3 kDa Amicon® concentrator. The absorbance at 280 nm was measured using a Nanodrop spectrophotometer to calculate sample concentration using the Beer-Lambert law [62] and extinction coefficients computed using the Expasy protparam website (**Supplementary Table 1**).

### RNA sample preparation

RNA sequences used in this study are included in **Supplementary Table 2**. Single-stranded RNA (ssRNA) oligonucleotides were purchased from IDT, resuspended in ultrapure water, and diluted to 1 mM in ultrapure water. 7SK SL3 hairpin RNA constructs were produced using *in vitro* RNA transcription with T7 RNA polymerase (Addgene #124138 [63], prepared in-house) and chemically synthesized DNA templates (IDT) following established protocols [18]. Briefly, T7 RNA Polymerase, 500 nM DNA template, and transcription buffer (40 mM Tris pH 8, 1 mM spermidine, 0.01% Triton-X, 40 mM MgCl_2_, 2.5 mM DTT, 20% DMSO, and 2 mM each rATP, rCTP, rUTP, rGTP) were incubated at 37 °C for 6-8 hours. RNA was purified by 15% denaturing polyacrylamide gel electrophoresis (PAGE) with 1x TBE running buffer and the RNA band was visualized by UV shadowing with a handheld UV lamp at 254 nm. After band excision, RNA was eluted from the gel using the ‘crush and soak’ method [64] by incubating gel pieces in crush and soak buffer (300 mM sodium acetate pH 5.2, 1 mM EDTA) for 24-48 hours at room temperature. RNA was further purified to remove acrylamide contaminants by ion-exchange chromatography using a diethylaminoethanol (DEAE) column (GE Healthcare) and elution into buffer (10 mM sodium phosphate pH 7.6, 1 mM EDTA, 1.5 M KCl). RNA was diluted to <100 μM in ultrapure water and annealed by heating to 95 °C for 3 min, followed by snap cooling on ice for 1 hour. RNA was then buffer exchanged into RNA sample buffer (20 mM sodium phosphate pH 6.0, 50 mM KCl, 0.1 mM EDTA) using a 3 kDa Amicon concentrator and concentrated to 0.8–1 mM.

### NMR spectroscopy

Solution NMR spectroscopy experiments were performed at 293.15 K on a Bruker Neo 600 MHz NMR spectrometer equipped with a triple-resonance cryoprobe. NMR samples were prepared in RBM7 storage buffer with added 5% D_2_O at 0.1-0.5 mM concentrations in 3 mm NMR tubes (Norell). Amide backbone resonance assignments were obtained from Biological Magnetic Resonance Data Bank (BMRB) [65] entry 19252. The Talos+ Server [66] was used to compute predicted *S*^*2*^ order parameter values using the BMRB-deposited STAR file as input. ^1^H-^15^N heteronuclear nuclear Overhauser effect (NOE) experiments (hsqcnoef3gpsi) from the Bruker experimental suite were recorded in an interleaved manner with 24 scans and 2 s incremental delay. The heteronuclear NOE is reported as the residue-specific ratio of peak intensity between the saturated and unsaturated experiments (I_sat_/I_unsat_). Error was estimated as the standard deviation of noise in the saturated *(σI*_*sat*_) and unsaturated (*σI*_*unsat*_) experiments [67, 68]:

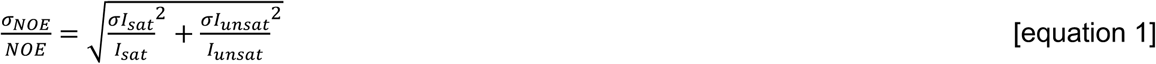

Data were processed using NMRPipe [69] and analyzed using NMRFAM-Sparky 1.470 powered by Sparky 3.190 [70] in the NMRbox virtual machine [71]. Chemical shift perturbation (CSP) experiments were performed using 0.2 mM protein in RBM7 storage buffer/5% D_2_O, with a reference spectrum recorded prior to addition of final concentrations of 0.02 mM (0.1 equivalent) and 0.1 mM (0.5 equivalent) RNA sequentially added to the NMR sample.

### iCLIP data analysis

RBM7 iCLIP sequencing data [34] were obtained from accession code E-MTAB-6475 in the EMBL-EBI database. Data were processed using the University of Nebraska-Lincoln Holland Computing Center (HCC) following the workflow reported in previous studies [72, 73]. First, FastQC [74] was run to analyze the per-base sequence quality of the data, per-base sequence content, barcode content and other quality content. Next, trim_galore [75] was used to trim the adapters and enhance the quality of the reads according to the quality obtained from the raw data. Next, iCount [76] was used to demultiplex the barcode sequences. Next, STAR [77] was used to index the GRChg38.p13 genome v34 (Gencode), align the reads to the genome, and generate a BAM file. Bedtools 2.27 [78] was used to convert the file of aligned reads from BAM to BED format, shift reads by one nt to account for premature reverse transcriptase (RT) termination adjacent to the crosslinked site (hereafter referred to as RT stop), extract the residue position number of the RT stop, and concatenate reads to map read coverage across the genome. The BED file was used as input for an in-house python program (https://github.com/ceichhorn2/7SKRNAextraction) to extract reads from RN7SK (ENSG00000202198, chr6: 52995620-52995950), convert to 7SK RNA residue number (1-331), and produce a file with the RT stop position number, the associated chromosome number, and the number of reads with the RT stop position number. The read counts for RT stops for a given residue (positional RT stop counts) were normalized to the total number of 7SK reads using the following equation:

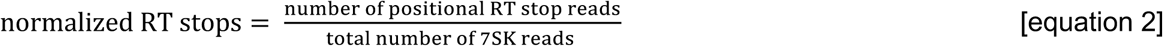

The average and standard deviation of the normalized RT stop values of the DMSO control (2 replicates) and 4-NQO treated (4 replicates) experiments were calculated using Excel (**Supplementary File 2**) and plots were generated using matplotlib. The average normalized RT stop values in the 4-NQO treated experiments were plotted on a secondary structure model of the upper stem of 7SK SL3 RNA (residues 210-264) using VARNA [79].

### Electrophoretic Mobility Shift Assays (EMSA)

RNA and protein samples were prepared as described above and diluted to 50 μM and 170 μM, respectively, in RBM7 storage buffer. To perform EMSA experiments, increasing amounts of protein samples were added to 50 pmol RNA (**Supplementary Table 3**). A protein-only sample was loaded in the first lane as a control to identify any nucleic acid contamination in the protein sample. The mixture was incubated on ice for 15-20 minutes, then 1 μL of loading dye (30% glycerol, bromophenol blue) was added for a total volume of 7.9 μL. Samples were loaded and run on 10% PAGE (37.5:1 bisacrylamide:acrylamide crosslinking ratio) at 50-60 volts for 40-80 minutes depending on RNA construct length. RNA was visualized with toluidine blue stain for hairpin and SYBR Safe (ThermoScientific) for ssRNA constructs.

## Results

### Extending the RBM7 RRM C-terminus improves expression and solubility

X-ray crystallographic and solution state NMR structures of the human RBM7 RRM have been previously determined using constructs with varying C-terminal boundaries [58, 59, 80]. For all solved structures, the RRM contains the canonical 1-α1-β2-β3-α2-β4 topology between amino acid (aa) residues T11-Q83; however, X-ray crystal structures report an additional β4’ strand in the α2-β4 loop and extended β4 strand. Constructs used for X-ray crystallography had minimal C-terminal ends ending at either position 86 [58] or 91 [59] whereas the construct used for the solution NMR structural ensemble had a minimal N-terminus and an extended C-terminus (aa 6-94) [56]. We first attempted to recombinantly express the full-length human RBM7 protein with an N-terminal His_6_ tag (HT-RBM7^FL^); however, this construct did not solubly express in *E. coli* (**Supplementary Figure S1**). We next generated constructs of RBM7 RRM with varying C-terminal ends: 1-86, 1-91, and 1-101 (named HT-RBM7^86^, HT-RBM7^91^, HT-RBM7^101^). To assess the impact of solubility tags on protein expression and solubility, additional fusion constructs were generated including either MBP or GB1 domains between His_6_ and TEV sites (HMT-RBM7^91^ and HGT-RBM7^101^, respectively) (**Fig. 1C**). Protein expression was greatest for HMT-RBM7^91^ and HGT-RBM7^101^ constructs (approximately 50 mg purified protein per L cell culture); however, despite successful TEV protease cleavage neither His_6_-MBP nor His_6_-GB1 could be separated from the RBM7 domain after TEV cleavage. Both MBP and GB1 are acidic (pI 5.1 and 4.5, respectively); it is likely that these domains have electrostatic interactions to the basic RBM7 (pI ∼9.4) (**Fig. 2A-B**).

**Figure 2.**
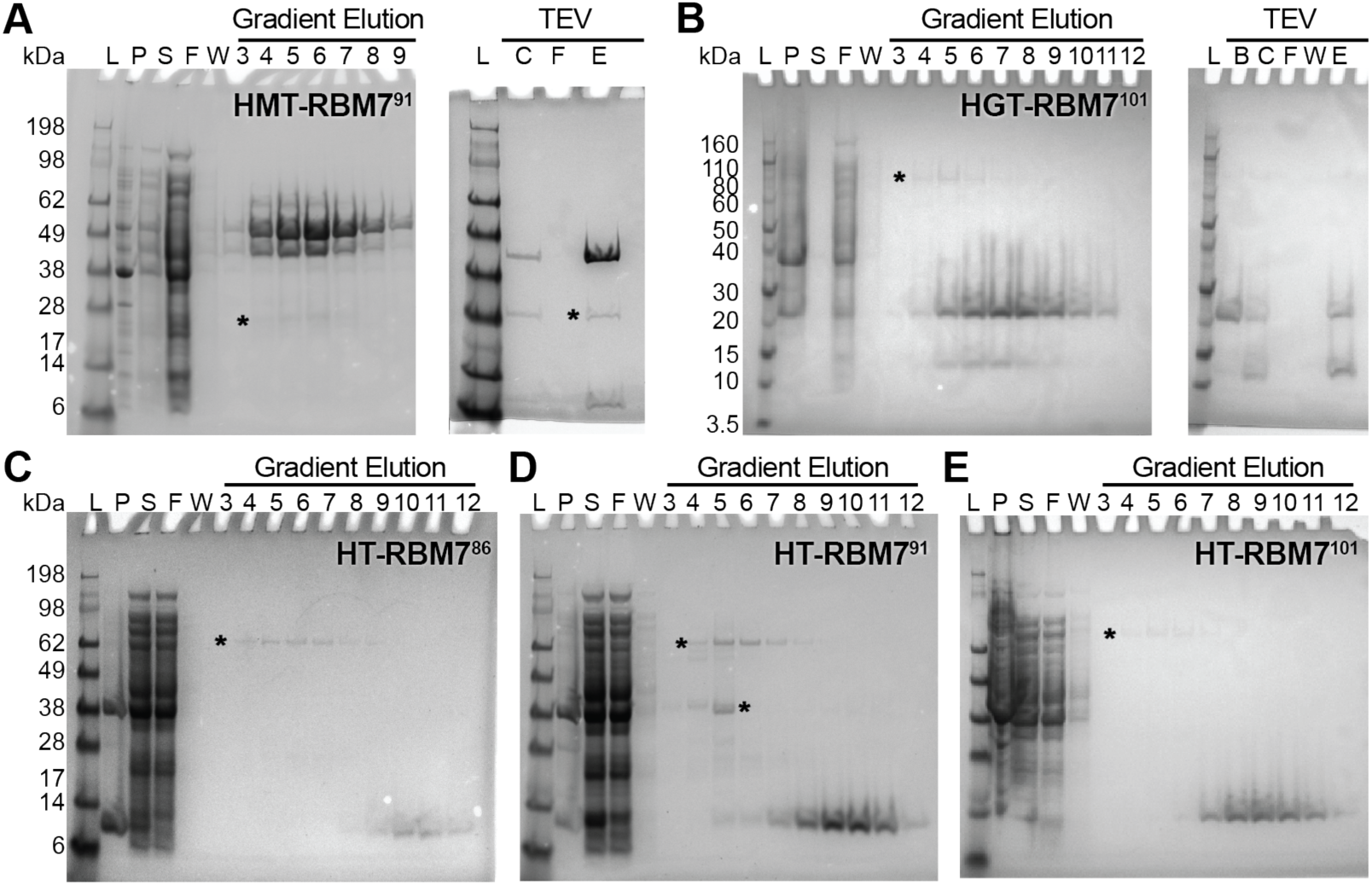
SDS-PAGE of recombinant protein expression and purification of RBM7 constructs. A-E *left*: L: molecular weight ladder; P: insoluble cell pellet after lysis; S: soluble fraction after lysis; F: flow-through from Ni-NTA column; W: wash from Ni-NTA column; 3-12: fraction number from Ni-NTA gradient elution. A, B *right)* B: sample before TEV cleavage; C: sample after TEV cleavage; F: flow-through from Ni-NTA column after TEV cleavage; W: wash from Ni-NTA column after TEV cleavage; E) elution from Ni-NTA column after TEV cleavage. SDS-PAGE of constructs with solubility tags A) MBP or B) GB1 showed high levels of soluble expression (*left*) and while TEV protease successfully cleaved RBM7 from the solubility tag RBM7 was retained on the Ni-NTA column (*right*). C) SDS-PAGE of constructs with His_6_ tag and varying C-terminal boundaries C) HT-RBM7^86^, D) HT-RBM7^91^, and E) HT-RBM7^101^. Asterisks indicate endogenous *E. coli* proteins that were observed as minor species that were successfully removed during Ni-NTA affinity purification.

The minimal HT-RBM7^86^ construct had poor yield after purification (approximately 2 mg per L cell culture), with significant amounts of protein observed in the insoluble fraction after cell lysis (**Fig. 2C**). HT-RBM7^86^ exhibited poor solubility and readily precipitated throughout purification and during concentration. Compared to HT-RBM7^86^, both HT-RBM7^91^ and HT-RBM7^101^ had higher yields (approximately 5-10 mg per L cell culture) and little protein observed in the insoluble fraction after cell lysis (**Fig. 2D-E**). Solution state NMR spectroscopy was used to evaluate HT-RBM7^86^, HT-RBM7^91^, and HT-RBM7^101^ construct folding and stability. The chemical shift assignments of the RBM7 RRM construct used to determine the solution NMR structural ensemble (aa 6-94) [56] were transferred onto 2D amide ^1^H-^15^N HSQC spectra, with good agreement between deposited and observed chemical shifts. The deposited chemical shifts lack assignments for the β3-α2 loop (aa K58-V63) and C-terminal residues (aa R85-S86, S89-H90). Peaks assigned to residues G87-D94 in the 2D amide ^1^H-^15^N HSQC spectrum of HT-RBM7^86^ were not observed, as expected. The spectrum showed reasonable chemical shift dispersion although several resonances corresponding to β1 (F13, G15, L17), α1 (Q32, G34), β2 (I37, K38), 3 (Q51-V55, F57), and β4 (I80, I82, Q83) suffered from extreme line-broadening, were absent, or shifted relative to constructs with extended C-termini indicative of global chemical exchange and conformational instability (**Fig. 3A**). Within 24 hours the sample began to precipitate, indicating poor solubility and stability of the minimal HT-RBM7^86^ construct.

**Figure 3.**
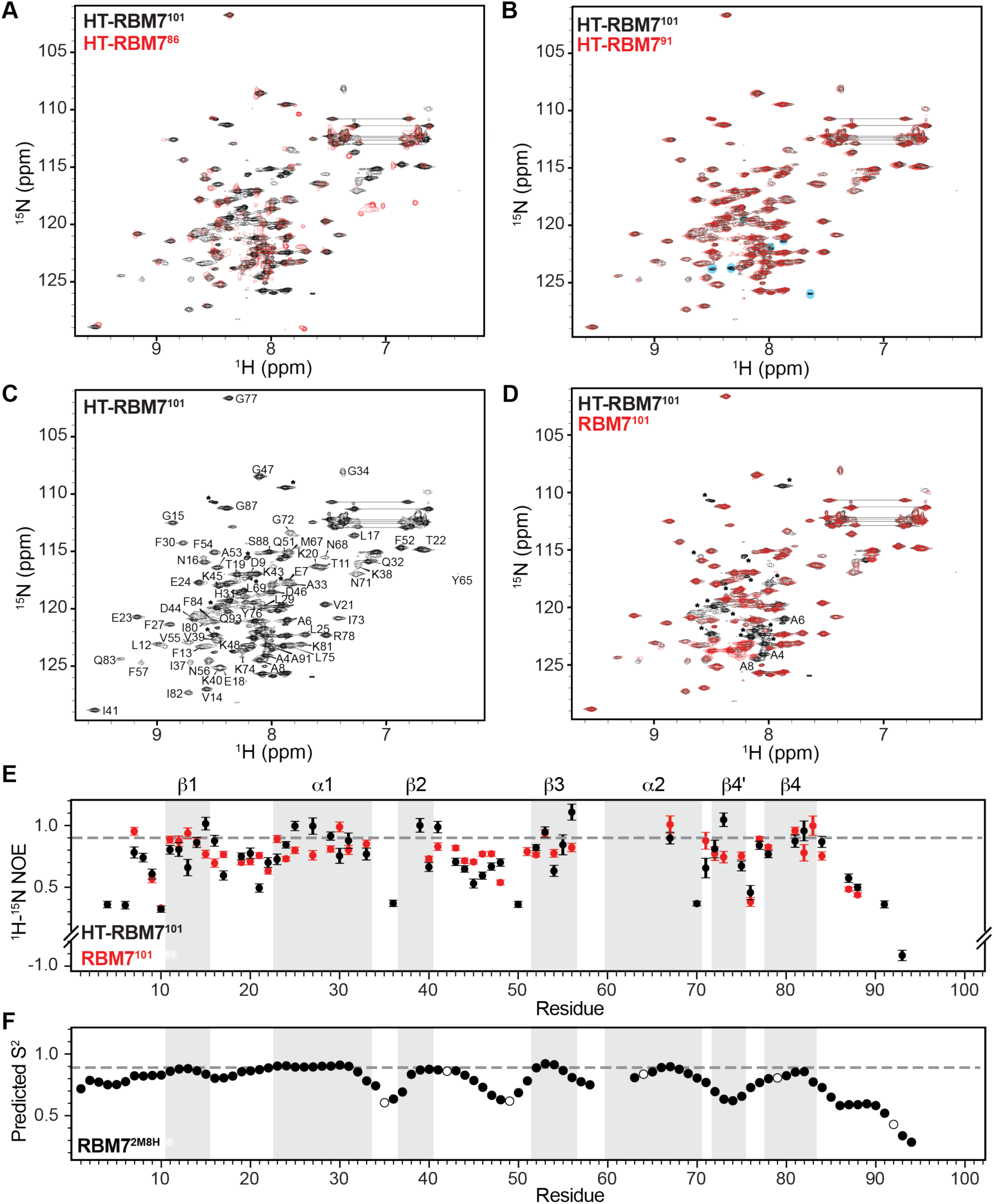
Comparison of RBM7 constructs by solution state NMR. Backbone amide 2D ^1^H-^15^N HSQCs of A) HT-RBM7^86^ (red) and B) HT-RBM7^91^ (red) overlaid with HT-RBM7^101^ (black). Blue circles indicate peaks observed in HT-RBM7^101^ and not HT-RBM7^91^. C) 2D ^1^H-^15^N HSQC of HT-RBM7^101^ with assignments transferred from BMRB entry 19252. D) ^1^H-^15^N HSQC spectra of RBM7^101^ (red) overlaid with HT-RBM7^101^ (black). Asterisks indicate peaks attributed to affinity tag, and labeled peaks are those assigned to the RRM that are missing or shifted. E) Plot of ^1^H-^15^N heteronuclear NOE values for HT-RBM7^101^(black) and RBM7^101^ (red). Gray box indicates secondary structure elements. F) Talos+ predicted S^2^ values from BMRB entry 19252. Proline residues are shown as open circles.

In contrast, the 2D amide ^1^H-^15^N HSQC spectrum of HT-RBM7^91^ showed excellent dispersion and uniform peak intensity (**Fig. 3B**). The majority of RRM resonances were observed and able to be assigned; however, the spectral quality was not stable over time and several resonances were no longer observed 4 days after sample preparation (**Fig. 3B and Supplementary Figure S2A**). These resonances correspond to the same missing resonances in HT-RBM7^86^ suggesting that while HT-RBM7^91^ is more stable, this construct experiences similar chemical exchange and construct instability. Similar to HT-RBM7^91^, HT-RBM7^101^ showed excellent chemical shift dispersion and uniform peak intensity (**Fig. 3C**). The ^1^H-^15^N HSQC spectra of HT-RBM7^91^ and HT-RBM7^101^ were nearly identical (**Fig. 3B**). We expected to observe ten additional resonances when comparing HT-RBM7^91^ and HT-RBM7^101^ and observed approximately six additional resonances in HT-RBM7^101^, of which several overlapped with other resonances (**Fig. 3B**). Unlike HT-RBM7^86^ and HT-RBM7^91^, the spectral quality of HT-RBM7^101^ remained consistent over time, with no loss of resonances seven days after sample purification (**Fig. 3C and Supplementary Figure S2B**).

### N-terminal His_6_ tag minimally impacts RRM stability and dynamics

Due to the observed interaction of the RBM7 RRM with solubility tags, we next evaluated the potential impact of the N-terminal His_6_ tag on RRM folding using solution NMR. The construct RBM7^101^ was generated by using TEV protease to cleave the His_6_ tag (**Fig. 3D**). The 2D amide ^1^H-^15^N HSQC spectra of HT-RBM7^101^ and RBM7^101^ are nearly identical for resonances assigned to the RRM (**Fig. 3D**). Approximately fifteen resonances are absent in the RBM7^101^ construct that are attributed to the sixteen N-terminal residues cleaved from HT-RBM7^101^. In addition, chemical shift perturbations are observed for N-terminal residues adjacent to the TEV cleavage site (A4, A6, A8) (**Fig. 3D**).

To assess RBM7 RRM conformational dynamics for both HT-RBM7^101^ and RBM7^101^ constructs, we performed ^1^H-^15^N heteronuclear NOE experiments, which report on ps-ns motional timescales (**Fig. 3E**). NOE values for several residues were unable to be determined due to lack of assignments particularly for α2 (K58-V63) and the C-terminus (D94-Q101), severe line broadening (F57, Y65, N68), and resonance overlap. The NOE values for HT-RBM7^101^ and RBM7^101^ are overall similar, indicating that the His_6_ tag does not alter RRM global folding and dynamics. Reduced values are observed for N- and C-terminal residues, indicating gradually increasing disorder at the termini. In addition, reduced values are observed in β1-α2 loop residues L17-E23, α1-β2 loop residue V36, β2-β3 loop residues K43-Q51, α2 residue L70, and α2-β4 loop residue Y76. V36 and Q50, which are adjacent to proline residues in the α1-β2 loop and β2-β3 loop respectively, have particularly reduced values. G72-K74 and R78-P79, which are observed to be α2-β4 loop residues in the NMR structure and β4’−β4 strands in X-ray structures, respectively (**Supplemental Figure S3**), have NOE values consistent with β-strand and α-helical residues in the RRM (**Fig. 3E**). Together, the heteronuclear NOE data shows a disordered N- and C-termini, flexible β2-β3 loop, and presence of a β4’ strand and extended β4 strand in both HT-RBM7^101^ and RBM7^101^ constructs.

In the prior study determining the solution NMR structural ensemble, the buffer conditions differed in pH and ionic strength; in addition, we found that inclusion of L-arginine and L-glutamic acid improved construct solubility. Further, the construct for structure determination had shortened N- and C-termini (aa 6-94) compared to our construct (aa 1-101), which together may account for the observed structural differences in the α2-β4 region. To further compare these two constructs, we used the chemical shift information in the BMRB deposition as input for the Talos+ server [66] to compute predicted secondary structure and *S*^*2*^ order parameters for the construct used previously for structural studies (RBM7^2M8H^) (**Fig. 3F**). The overall pattern of the predicted *S*^*2*^ values for RBM7^2M8H^ shows excellent agreement to our observed NOE values, particularly proline-adjacent residues V36 and Q50 and C-terminal residues. The predicted *S*^*2*^ values show decreased values for residues G72-Y76 and the secondary structure of these residues is predicted to be unstructured, supporting the determined solution NMR structure model in which these residues are disordered.

### RBM7 RRM binds 7SK RNA stem-loop 3

RBM7 was previously identified to interact with the 7SK RNP using iCLIP experiments that showed RBM7 crosslinking to the stem-loop 3 (SL3) region of 7SK RNA; however, the precise binding site was not defined in the prior study [34]. In addition, mutagenesis of the RRM domain showed that the RBM7 RRM RNP1 and RNP2 sequences were required for 7SK RNP association *in cellulo* [34]. To gain additional insights into specific 7SK RNA residues that crosslinked to RBM7, we obtained iCLIP data deposited to EMBL and processed and analyzed data. The per-residue 7SK RT stops were normalized to the total number of 7SK reads for both DMSO control and 4-NQO treated samples to measure crosslinking enrichment across the 7SK RNA sequence (**Fig. 4A**). Consistent with the previous study, RBM7 crosslinking to 7SK RNA was observed in both control and treated samples (**Fig. 4A**) [34] suggesting an interaction between RBM7 and the 7SK RNP even in the absence of DNA damage. Although the 7SK SL3 secondary structure and boundaries are not well characterized, independent chemical mapping studies report SL3 boundaries between residues 200-274 [81-83]. Crosslinking was enriched in the upper stem of 7SK SL3 and observed to be highest for residues A245-U246, with additional crosslinking observed adjacent to this site as well as G232-C235 (**Fig. 4B**).

**Figure 4.**
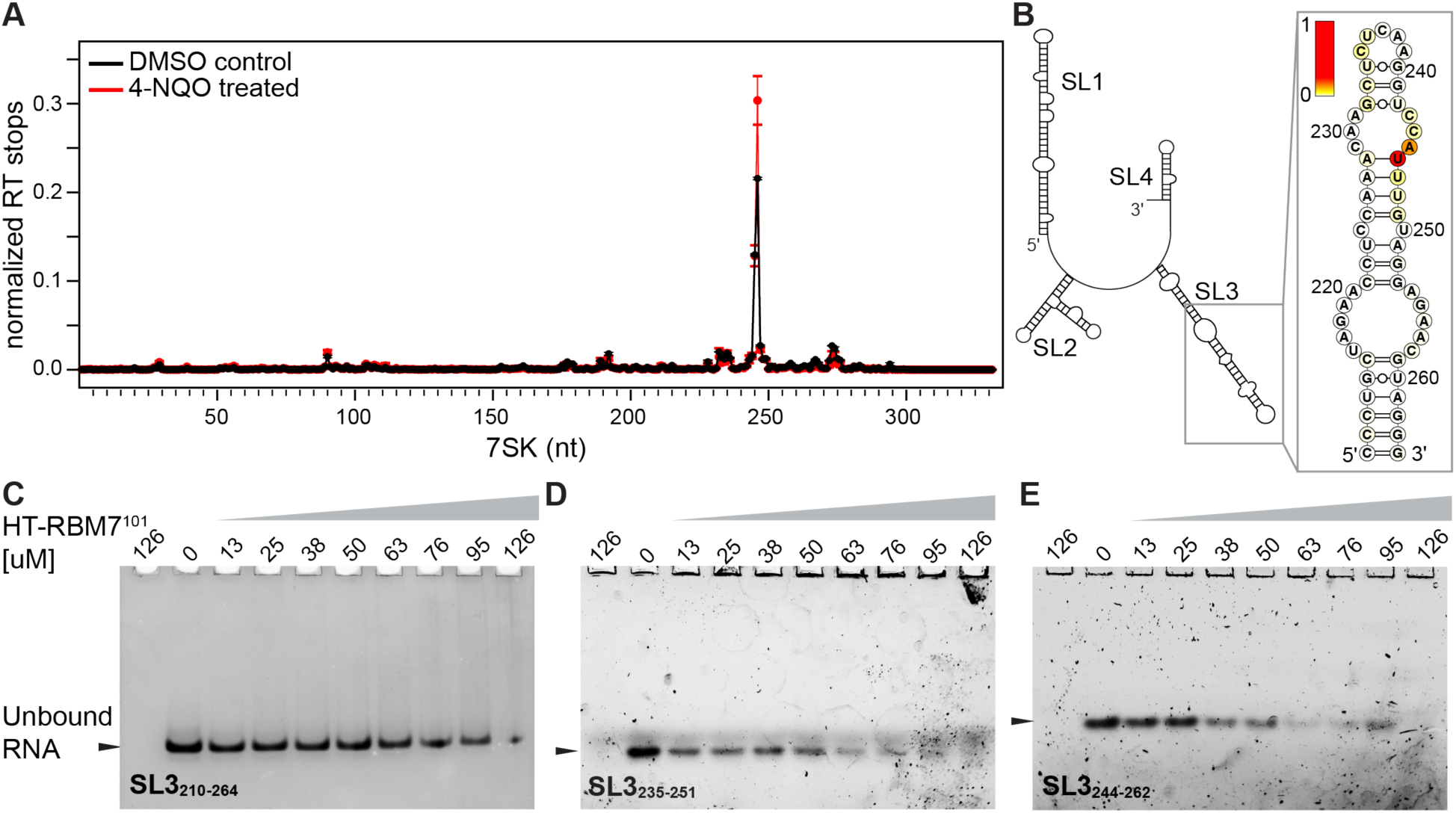
RBM7 interaction with 7SK RNA. A) Plot of average normalized RT stop values from DMSO control (black) and 4-NQO-treated (red) iCLIP experiments. B) Average normalized RT stop values of 4-NQO-treated sample, shown in panel A, plotted as a heat map onto a secondary structure model of the upper region of 7SK SL3. C-E) EMSA of HT-RBM7^101^ with 7SK SL3 RNA constructs for C) hairpin SL3_210-264_, D) ssRNA SL3_235-251_, E) ssRNA SL3_244-262_.

To investigate RBM7 RRM binding to 7SK SL3 *in vitro*, we performed qualitative EMSAs using HT-RBM7^101^ and various constructs of 7SK SL3 comprising either hairpin or ssRNA sequences (**Fig. 4C-F**). To ensure protein samples did not contain contaminating nucleic acids that co-purified from *E. coli*, a control lane containing only protein was included. Unfortunately, we observed sample precipitation shortly after addition of protein to RNA, with EMSAs showing only unbound RNA and not the corresponding protein-bound band. Binding was inferred from the decreasing intensity of the unbound band with increasing protein concentration. We first used a hairpin construct comprising the upper stem of 7SK RNA SL3 (SL3_210-264_) (**Fig. 4B**) and observed extremely weak binding requiring significant HT-RBM7^101^ stoichiometric excess (**Fig. 4C**). Given that canonical RRMs generally bind ssRNA [84], and the RBM7 RRM preferentially binds to pyrimidine-rich ssRNA [56] we then assessed the binding of ssRNA constructs corresponding to 7SK SL3 regions with enriched crosslinking to RBM7: SL3_235-251_, which includes the SL3 apical loop, and SL3_244-262_ (**Fig. 4D-E**). Although these constructs also require excess protein to saturate binding, both constructs show relatively improved binding compared to the hairpin construct.

We performed solution NMR chemical shift perturbation (CSP) experiments to probe binding of HT-RBM7^101^ to SL3_244-262_. Similar to EMSA experiments, immediate precipitation of the protein was observed upon addition of RNA. However, we were able to collect ^1^H-^15^N HSQC spectra of HT-RBM7^101^ with 0.1 and 0.5 equivalents of SL3_244-262_ (**Fig. 5**). Addition of 0.1 eq RNA uniformly reduced the resonance signal intensity compared to protein in the absence of RNA, indicating that the protein signal loss is due to RNA-protein complex formation and subsequent precipitation. The ^1^H-^15^N HSQC spectra of HT-RBM7^101^ in the absence and presence of 0.1 eq RNA is identical with no apparent chemical shift perturbations, suggesting that the observed resonances are unbound protein. At 0.5 eq, the majority of RRM resonances disappear, with only resonances of the His_6_ tag, N-terminus (A4, L12), and β1-α1 loop (V21, T22, E23, E24) observed (**Fig. 5**).

**Figure 5.**
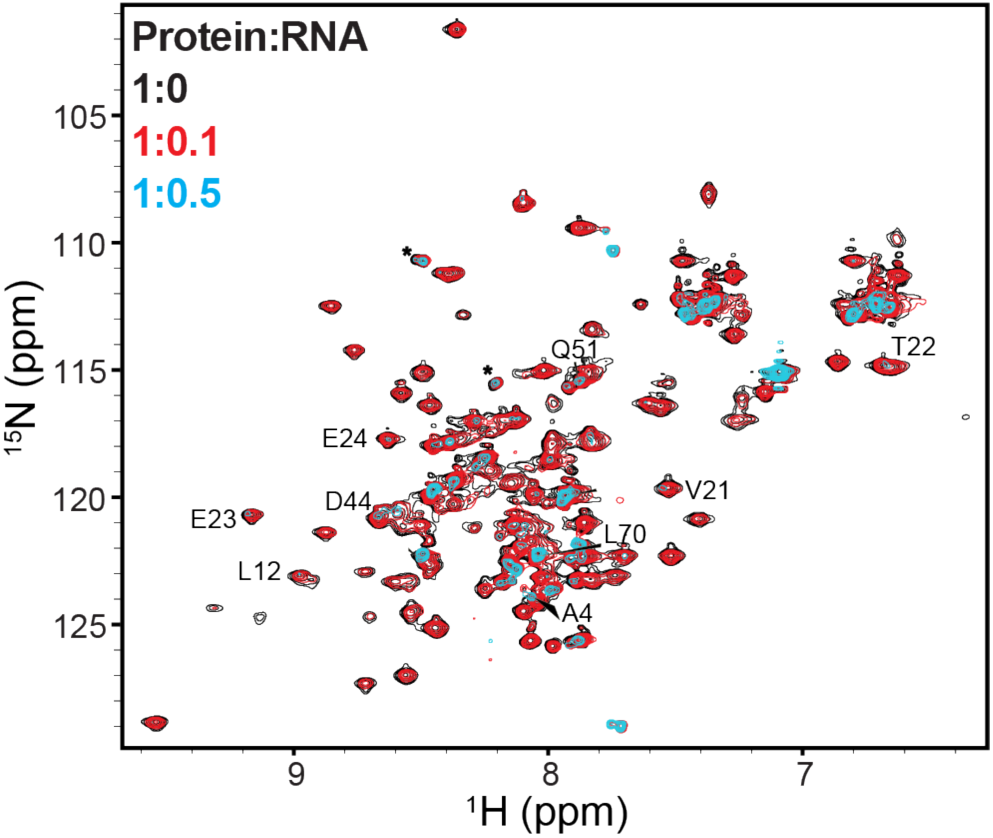
Chemical shift perturbation of HT-RBM7^101^ with SL3_244-262_ shows disappearance of free RBM7 with increasing RNA. HT-RBM7^101^ in absence of RNA shown in black, with 0.1 equivalents of SL3_244-262_ shown in red, and with 0.5 equivalents of SL3_244-262_ shown in blue. Labeled resonances are those that remain visible with addition of 0.5 equivalents RNA.

## Discussion

As a component of NEXT, RBM7 targets RNA for processing and turnover [53, 54]. An alternate function for RBM7 has recently been identified, independent of NEXT, in which RBM7 assembles with 7SK RNP to promote P-TEFb release and subsequent transcription activation of genes in response to genotoxic stress [34]. RBM7 assembly with proteins is essential for its function in NEXT [50, 58]. Similarly, RBM7 interacts with both 7SK RNA and core 7SK RNP protein components [34]. X-ray structures of the RBM7 RRM in complex with NEXT component ZCCHC8 (PDB IDs 5LXR, 5LXY) [58] show the RRM−ZCCHC8 interface to be on the underside of the RRM (α1-α2-β4’), opposite to the putative RNA-binding interface on the β-sheet surface (**Supplemental Figure S3**). Similarly, in the X-ray structure of the individual RBM7 RRM (PDB ID 5IQQ), the construct crystallized as a pentamer with observed intermolecular interactions at the α2-β4’-β4 interface (E60, P64, N68, K74, Y76, F84, R85) (**Supplemental Figure S4**). These observed crystal contacts may correspond to protein-protein interaction sites in solution, and the RBM7 RRM has previously been described to be prone to aggregation *in vitro* [80]. In further support of protein-protein interactions in this region of the RRM, NMR chemical shift assignments were not reported for residues K58-V63 in the solution NMR structure (PDB ID 2M8H) [56]. In this study, ^1^H-^15^N HSQC spectra show line-broadening of resonances for residues adjacent to this region (F57 in β3-α2 loop, Y65 and N68 in α2, N71 in α2-β4’ loop, G72 in β4’), indicating chemical exchange that may be due to transient protein-protein interactions. We also observe protein-protein interactions between acidic solubility tags and the RRM, likely governed by electrostatic interactions.

To investigate the C-terminal boundary requirements for RBM7 RRM folding and stability, we generated RRM constructs corresponding to the X-ray structure of the individual RRM (aa 1-86, PDB ID 5IQQ), RRM-ZCCHC8 complex (aa 1-91, PDB IDs 5LXR and 5LXY), and an extended construct ending at residue 101. We found that while all constructs could be recombinantly expressed and purified, the minimal HT-RBM7^86^ construct was the least stable with the lowest yields, rapid precipitation, and deterioration of ^1^H-^15^N HSQC spectral quality within one day. Although HT-RBM7^91^ showed improved solubility, this construct also showed similar deterioration of ^1^H-^15^N HSQC spectral quality within four days. The extended HT-RBM7^101^ construct showed the greatest stability, with no observed precipitation nor deterioration of NMR ^1^H-^15^N HSQC spectral quality. Constructs with shortened C-terminal ends may destabilize β4 strand positioning on the β-sheet, resulting in reduced thermodynamic stability of the RRM. Alternately, these minimal constructs may be more susceptible to protein-protein interactions and subsequent aggregation; the HT-RBM7^101^ extended C-terminus may prevent protein-protein interactions through steric hindrance. Addition of protein cofactors may further improve RBM7 RRM solubility.

For all determined structures, the N- and C-termini are disordered, with either missing density [59] or high conformational variability in the solution NMR structural ensemble [56]. In addition, there is a discrepancy between the observed secondary structures in the solution NMR and X-ray structures, in which the X-ray structures contain an additional β4’ strand between α2 and β4, as well as an extended β4 strand relative to the solution NMR structure (**Supplemental Figure S3**). X-ray structures report a lowest-energy conformation whereas NMR structures report a structural ensemble that is often more conformationally dynamic compared to a crystal structure. The α2-β4region is part of the protein-protein interface (**Supplemental Figures 3-4**), and the β4’-extended β4 strands observed in the X-ray structures may be further stabilized by protein-protein interactions. Our heteronuclear NOE data shows that in the HT-RBM7^101^ construct these residues are ordered, consistent with β4’ and extended β4 strand formation. However, the Talos+ order parameters and secondary structure predicted from the chemical shifts of RBM7^2M8H^ suggest an unstructured loop rather than a β strand, in support of the solution NMR structure previously determined. Our study uses an extended construct, with additional residues at both the N- and C-termini. In addition, we use a different buffer that includes a reduced pH and amino acids L-arginine and L-glutamic acid to improve solubility. These differences in construct boundaries and buffer may account for the observed differences in secondary structure. Unusually, the beginning of the extended β strand contains a proline (P79). Prolines are generally disfavored in β-strands due to lacking an amide proton that participates in hydrogen bonding [85], and may reduce stable formation of the extended β4 strand, contributing to instability in this region.

Previously, the RBM7 RRM was shown to bind 7-mer ssRNA constructs, with affinities ranging from 60-150 μM [56]. Highest affinities were observed to polyU and alternating CU-repeat ssRNA sequences, with weak to no binding detected to polyC and polyA ssRNA sequences. The previous iCLIP study showed enriched crosslinking of RBM7 to 7SK SL3 [34], and in this study we localized RBM7-7SK RNA crosslinking to the distal region of 7SK SL3, which is enriched in unpaired pyrimidines. In addition, base pairs in the SL3 upper stem are primarily G·U and A−U base pairs, which are less thermodynamically stable compared to a G•C base pair. We show that *in vitro*, the RBM7 RRM has little observed binding to the hairpin 7SK SL3 RNA but binds to ssRNA constructs corresponding to the 7SK SL3 apical loop and upper stem. The relative increased binding of HT-RBM7^101^ to ssRNA compared to hairpin constructs suggests a mode of recognition in which RBM7 binds to unpaired residues in the distal region of 7SK SL3. In addition to 7SK RNA, RBM7 interacts with core 7SK RNP proteins MePCE and Larp7 [34]. Given the apparent weak binding of RBM7 to 7SK RNA, RBM7 assembly onto 7SK RNP likely requires both protein-protein and protein-RNA interactions for stable association with 7SK RNP and P-TEFb release. 7SK SL3 was recently shown to interact with another RNA binding protein, hnRNPA1 [82], which is also involved in P-TEFb release from 7SK RNP [86] and has increased association with 7SK RNP after 4-NQO treatment [34]. RBM7, hnRNPA1, and additional accessory proteins may work cooperatively with core 7SK RNP proteins to coordinate P-TEFb release for targeted regulation of specific genes in response to genotoxic stress.

## Data availability

Samples, reagents, and raw data files generated in this work are available upon reasonable request. In-house programs generated in this work are deposited on the GitHub repository https://github.com/ceichhorn2/7SKRNAextraction.

## Declaration of competing interest

The authors declare no competing interest.

## Acknowledgements

We acknowledge funding support from the National Institutes of Health (1R35GM143030), the Nebraska Center for Integrated Biomolecular Communication (P20 GM113126), and UNL startup funds to CDE and UNL UCARE funding to AMS. We thank Ernest Atsrim for critical review of the manuscript, Momodou Camara for assistance with RNA sample preparation, Youssef Fathy for assistance with programming, Dr. Martha Morton for NMR assistance, and Drs Chia-sin Liu and John-Jack Riethoven at the Holland Computing Center for assistance with processing iCLIP data. We gratefully acknowledge Addgene for plasmids used for plasmid generation and to prepare enzymes involved in preparation of RBM7 and *in vitro* transcribed RNA samples. pDZ2087 was a gift from David Waugh (Addgene plasmid #92414); pQE30-His-T7RNAP was a gift from Sebastian Maerkl & Takuya Ued (Addgene plasmid #124138); pET His6 MBP TEV LIC cloning vector (1M) was a gift from Scott Gradia (Addgene plasmid #29656).

## Notes

### Competing Interest Statement

The authors have declared no competing interest.

## References

[1] K. Adelman, J.T. Lis, Promoter-proximal pausing of RNA polymerase II: emerging roles in metazoans, Nat Rev Genet, 13 (2012) 720–731.

[2] Q. Zhou, T. Li, D.H. Price, RNA polymerase II elongation control, Annu Rev Biochem, 81 (2012) 119–143.

[3] I. Jonkers, J.T. Lis, Getting up to speed with transcription elongation by RNA polymerase II, Nat Rev Mol Cell Biol, 16 (2015) 167–177.

[4] L. Core, K. Adelman, Promoter-proximal pausing of RNA polymerase II: a nexus of gene regulation, Genes Dev, 33 (2019) 960–982.

[5] C.Q. Aj, A. Bugai, M. Barboric, Cracking the control of RNA polymerase II elongation by 7SK snRNP and P-TEFb, Nucleic Acids Res, 44 (2016) 7527–7539.

[6] R.P. McNamara, C.W. Bacon, I. D’Orso, Transcription elongation control by the 7SK snRNP complex: Releasing the pause, Cell Cycle, 15 (2016) 2115–2123.

[7] D. Ivanov, Y.T. Kwak, J. Guo, R.B. Gaynor, Domains in the SPT5 protein that modulate its transcriptional regulatory properties, Mol Cell Biol, 20 (2000) 2970–2983.

[8] J. Kohoutek, P-TEFb-the final frontier, Cell Div, 4 (2009) 19.

[9] N.F. Marshall, J. Peng, Z. Xie, D.H. Price, Control of RNA polymerase II elongation potential by a novel carboxyl-terminal domain kinase, J Biol Chem, 271 (1996) 27176–27183.

[10] Z. Yang, Q. Zhu, K. Luo, Q. Zhou, The 7SK small nuclear RNA inhibits the CDK9/cyclin T1 kinase to control transcription, Nature, 414 (2001) 317–322.

[11] V.T. Nguyen, T. Kiss, A.A. Michels, O. Bensaude, 7SK small nuclear RNA binds to and inhibits the activity of CDK9/cyclin T complexes, Nature, 414 (2001) 322–325.

[12] E. Ullu, V. Esposito, M. Melli, Evolutionary conservation of the human 7 S RNA sequences, J Mol Biol, 161 (1982) 195–201.

[13] Y. Xue, Z. Yang, R. Chen, Q. Zhou, A capping-independent function of MePCE in stabilizing 7SK snRNA and facilitating the assembly of 7SK snRNP, Nucleic Acids Res, 38 (2010) 360–369.

[14] Y. Yang, C.D. Eichhorn, Y. Wang, D. Cascio, J. Feigon, Structural basis of 7SK RNA 5’-gamma-phosphate methylation and retention by MePCE, Nat Chem Biol, 15 (2019) 132–140.

[15] B.J. Krueger, C. Jeronimo, B.B. Roy, A. Bouchard, C. Barrandon, S.A. Byers, C.E. Searcey, J.J. Cooper, O. Bensaude, E.A. Cohen, B. Coulombe, D.H. Price, LARP7 is a stable component of the 7SK snRNP while P-TEFb, HEXIM1 and hnRNP A1 are reversibly associated, Nucleic Acids Res, 36 (2008) 2219–2229.

[16] A. Markert, M. Grimm, J. Martinez, J. Wiesner, A. Meyerhans, O. Meyuhas, A. Sickmann, U. Fischer, The La-related protein LARP7 is a component of the 7SK ribonucleoprotein and affects transcription of cellular and viral polymerase II genes, EMBO Rep, 9 (2008) 569–575.

[17] E. Uchikawa, K.S. Natchiar, X. Han, F. Proux, P. Roblin, E. Zhang, A. Durand, B.P. Klaholz, A.-C. Dock-Bregeon, Structural insight into the mechanism of stabilization of the 7SK small nuclear RNA by LARP7, Nucleic Acids Res, 43 (2015) 3373–3388.

[18] C.D. Eichhorn, Y. Yang, L. Repeta, J. Feigon, Structural basis for recognition of human 7SK long noncoding RNA by the La-related protein Larp7, Proc Natl Acad Sci U S A, 115 (2018) E6457–E6466.

[19] Y. Yang, S. Liu, S. Egloff, C.D. Eichhorn, T. Hadjian, J. Zhen, T. Kiss, Z.H. Zhou, J. Feigon, Structural basis of RNA conformational switching in the transcriptional regulator 7SK RNP, Mol Cell, 82 (2022) 1724–1736 e1727.

[20] J.H. Yik, R. Chen, R. Nishimura, J.L. Jennings, A.J. Link, Q. Zhou, Inhibition of P-TEFb (CDK9/Cyclin T) kinase and RNA polymerase II transcription by the coordinated actions of HEXIM1 and 7SK snRNA, Mol Cell, 12 (2003) 971–982.

[21] A.A. Michels, A. Fraldi, Q. Li, T.E. Adamson, F. Bonnet, V.T. Nguyen, S.C. Sedore, J.P. Price, D.H. Price, L. Lania, O. Bensaude, Binding of the 7SK snRNA turns the HEXIM1 protein into a P-TEFb (CDK9/cyclin T) inhibitor, Embo J, 23 (2004) 2608–2619.

[22] L. Kobbi, E. Demey-Thomas, F. Braye, F. Proux, O. Kolesnikova, J. Vinh, A. Poterszman, O. Bensaude, An evolutionary conserved Hexim1 peptide binds to the Cdk9 catalytic site to inhibit P-TEFb, Proc Natl Acad Sci U S A, 113 (2016) 12721–12726.

[23] S. Egloff, E. Van Herreweghe, T. Kiss, Regulation of polymerase II transcription by 7SK snRNA: two distinct RNA elements direct P-TEFb and HEXIM1 binding, Mol Cell Biol, 26 (2006) 630–642.

[24] N. He, N.S. Jahchan, E. Hong, Q. Li, M.A. Bayfield, R.J. Maraia, K. Luo, Q. Zhou, A La-related protein modulates 7SK snRNP integrity to suppress P-TEFb-dependent transcriptional elongation and tumorigenesis, Mol Cell, 29 (2008) 588–599.

[25] M. Briese, M. Sendtner, Keeping the balance: The noncoding RNA 7SK as a master regulator for neuron development and function, Bioessays, 43 (2021) e2100092.

[26] P. Liu, Y. Xiang, K. Fujinaga, K. Bartholomeeusen, K.A. Nilson, D.H. Price, B.M. Peterlin, Release of positive transcription elongation factor b (P-TEFb) from 7SK small nuclear ribonucleoprotein (snRNP) activates hexamethylene bisacetamide-inducible protein (HEXIM1) transcription, J Biol Chem, 289 (2014) 9918–9925.

[27] J.R. Hogg, K. Collins, RNA-based affinity purification reveals 7SK RNPs with distinct composition and regulation, RNA, 13 (2007) 868–880.

[28] T. Cherrier, V. Le Douce, S. Eilebrecht, R. Riclet, C. Marban, F. Dequiedt, Y. Goumon, J.C. Paillart, M. Mericskay, A. Parlakian, P. Bausero, W. Abbas, G. Herbein, S.K. Kurdistani, X. Grana, B. Van Driessche, C. Schwartz, E. Candolfi, A.G. Benecke, C. Van Lint, O. Rohr, CTIP2 is a negative regulator of P-TEFb, Proc Natl Acad Sci U S A, 110 (2013) 12655–12660.

[29] S. Eilebrecht, B.J. Benecke, A. Benecke, 7SK snRNA-mediated, gene-specific cooperativity of HMGA1 and P-TEFb, RNA Biol, 8 (2011) 1084–1093.

[30] S. Eilebrecht, E. Wilhelm, B.J. Benecke, B. Bell, A.G. Benecke, HMGA1 directly interacts with TAR to modulate basal and Tat-dependent HIV transcription, RNA Biol, 10 (2013) 436–444.

[31] R.P. McNamara, J.E. Reeder, E.A. McMillan, C.W. Bacon, J.L. McCann, I. D’Orso, KAP1 Recruitment of the 7SK snRNP Complex to Promoters Enables Transcription Elongation by RNA Polymerase II, Mol Cell, 61 (2016) 39–53.

[32] S.A. Gudipaty, R.P. McNamara, E.L. Morton, I. D’Orso, PPM1G Binds 7SK RNA and Hexim1 To Block P-TEFb Assembly into the 7SK snRNP and Sustain Transcription Elongation, Mol Cell Biol, 35 (2015) 3810–3828.

[33] E. Calo, R.A. Flynn, L. Martin, R.C. Spitale, H.Y. Chang, J. Wysocka, RNA helicase DDX21 coordinates transcription and ribosomal RNA processing, Nature, 518 (2015) 249–253.

[34] A. Bugai, A.J.C. Quaresma, C.C. Friedel, T. Lenasi, R. Duster, C.R. Sibley, K. Fujinaga, P. Kukanja, T. Hennig, M. Blasius, M. Geyer, J. Ule, L. Dolken, M. Barboric, P-TEFb Activation by RBM7 Shapes a Pro-survival Transcriptional Response to Genotoxic Stress, Mol Cell, 74 (2019) 254–267 e210.

[35] M.K. Jang, K. Mochizuki, M. Zhou, H.S. Jeong, J.N. Brady, K. Ozato, The bromodomain protein Brd4 is a positive regulatory component of P-TEFb and stimulates RNA polymerase II-dependent transcription, Mol Cell, 19 (2005) 523–534.

[36] Q. Zhou, J.H. Yik, The Yin and Yang of P-TEFb regulation: implications for human immunodeficiency virus gene expression and global control of cell growth and differentiation, Microbiol Mol Biol Rev, 70 (2006) 646–659.

[37] C.W. Bacon, I. D’Orso, CDK9: a signaling hub for transcriptional control, Transcription, 10 (2019) 57–75.

[38] M. Barboric, T. Lenasi, Kick-sTARting HIV-1 transcription elongation by 7SK snRNP deporTATion, Nat Struct Mol Biol, 17 (2010) 928–930.

[39] M. Bouhaddou, D. Memon, B. Meyer, K.M. White, V.V. Rezelj, M. Correa Marrero, B.J. Polacco, J.E. Melnyk, S. Ulferts, R.M. Kaake, J. Batra, A.L. Richards, E. Stevenson, D.E. Gordon, A. Rojc, K. Obernier, J.M. Fabius, M. Soucheray, L. Miorin, E. Moreno, C. Koh, Q.D. Tran, A. Hardy, R. Robinot, T. Vallet, B.E. Nilsson-Payant, C. Hernandez-Armenta, A. Dunham, S. Weigang, J. Knerr, M. Modak, D. Quintero, Y. Zhou, A. Dugourd, A. Valdeolivas, T. Patil, Q. Li, R. Huttenhain, M. Cakir, M. Muralidharan, M. Kim, G. Jang, B. Tutuncuoglu, J. Hiatt, J.Z. Guo, J. Xu, S. Bouhaddou, C.J.P. Mathy, A. Gaulton, E.J. Manners, E. Felix, Y. Shi, M. Goff, J.K. Lim, T. McBride, M.C. O’Neal, Y. Cai, J.C.J. Chang, D.J. Broadhurst, S. Klippsten, E. De Wit, A.R. Leach, T. Kortemme, B. Shoichet, M. Ott, J. Saez-Rodriguez, B.R. tenOever, R.D. Mullins, E.R. Fischer, G. Kochs, R. Grosse, A. Garcia-Sastre, M. Vignuzzi, J.R. Johnson, K.M. Shokat, D.L. Swaney, P. Beltrao, N.J. Krogan, The Global Phosphorylation Landscape of SARS-CoV-2 Infection, Cell, 182 (2020) 685–712 e619.

[40] X. Ji, Y. Zhou, S. Pandit, J. Huang, H. Li, C.Y. Lin, R. Xiao, C.B. Burge, X.D. Fu, SR proteins collaborate with 7SK and promoter-associated nascent RNA to release paused polymerase, Cell, 153 (2013) 855–868.

[41] R.P. McNamara, C. Guzman, J.E. Reeder, I. D’Orso, Genome-wide analysis of KAP1, the 7SK snRNP complex, and RNA polymerase II, Genom Data, 7 (2016) 250–255.

[42] M.D. Lavigne, D. Konstantopoulos, K.Z. Ntakou-Zamplara, A. Liakos, M. Fousteri, Global unleashing of transcription elongation waves in response to genotoxic stress restricts somatic mutation rate, Nat Commun, 8 (2017) 2076.

[43] C. Studniarek, M. Tellier, P.G.P. Martin, S. Murphy, T. Kiss, S. Egloff, The 7SK/P-TEFb snRNP controls ultraviolet radiation-induced transcriptional reprogramming, Cell Rep, 35 (2021) 108965.

[44] M. Blasius, S.A. Wagner, C. Choudhary, J. Bartek, S.P. Jackson, A quantitative 14-3-3 interaction screen connects the nuclear exosome targeting complex to the DNA damage response, Genes Dev, 28 (2014) 1977–1982.

[45] M.E. Borisova, A. Voigt, M.A.X. Tollenaere, S.K. Sahu, T. Juretschke, N. Kreim, N. Mailand, C. Choudhary, S. Bekker-Jensen, M. Akutsu, S.A. Wagner, P. Beli, p38-MK2 signaling axis regulates RNA metabolism after UV-light-induced DNA damage, Nat Commun, 9 (2018) 1017.

[46] H.C. Reinhardt, A.S. Aslanian, J.A. Lees, M.B. Yaffe, p53-deficient cells rely on ATM-and ATR-mediated checkpoint signaling through the p38MAPK/MK2 pathway for survival after DNA damage, Cancer Cell, 11 (2007) 175–189.

[47] H. Sobhy, R. Kumar, J. Lewerentz, L. Lizana, P. Stenberg, Highly interacting regions of the human genome are enriched with enhancers and bound by DNA repair proteins, Sci Rep, 9 (2019) 4577.

[48] S.T. Younger, J.L. Rinn, p53 regulates enhancer accessibility and activity in response to DNA damage, Nucleic Acids Res, 45 (2017) 9889–9900.

[49] C.A. Higgins, B.E. Nilsson-Payant, A.P. Kurland, P. Adhikary, I. Golynker, O. Danziger, M. Panis, B.R. Rosenberg, B. tenOever, J.R. Johnson, SARS-CoV-2 hijacks p38ß/MAPK11 to promote viral protein translation, bioRxiv, (2021) 2021.2008.2020.457146.

[50] M.P. Gustafson, M. Welcker, H.C. Hwang, B.E. Clurman, Zcchc8 is a glycogen synthase kinase-3 substrate that interacts with RNA-binding proteins, Biochem Biophys Res Commun, 338 (2005) 1359–1367.

[51] M. Lubas, M.S. Christensen, M.S. Kristiansen, M. Domanski, L.G. Falkenby, S. Lykke-Andersen, J.S. Andersen, A. Dziembowski, T.H. Jensen, Interaction profiling identifies the human nuclear exosome targeting complex, Mol Cell, 43 (2011) 624–637.

[52] M. Hallais, F. Pontvianne, P.R. Andersen, M. Clerici, D. Lener, H. Benbahouche Nel, T. Gostan, F. Vandermoere, M.C. Robert, S. Cusack, C. Verheggen, T.H. Jensen, E. Bertrand, CBC-ARS2 stimulates 3’-end maturation of multiple RNA families and favors cap-proximal processing, Nat Struct Mol Biol, 20 (2013) 1358–1366.

[53] M. Lubas, P.R. Andersen, A. Schein, A. Dziembowski, G. Kudla, T.H. Jensen, The human nuclear exosome targeting complex is loaded onto newly synthesized RNA to direct early ribonucleolysis, Cell Rep, 10 (2015) 178–192.

[54] C. Kilchert, S. Wittmann, L. Vasiljeva, The regulation and functions of the nuclear RNA exosome complex, Nat Rev Mol Cell Biol, 17 (2016) 227–239.

[55] C. Tiedje, M. Lubas, M. Tehrani, M.B. Menon, N. Ronkina, S. Rousseau, P. Cohen, A. Kotlyarov, M. Gaestel, p38MAPK/MK2-mediated phosphorylation of RBM7 regulates the human nuclear exosome targeting complex, RNA, 21 (2015) 262–278.

[56] D. Hrossova, T. Sikorsky, D. Potesil, M. Bartosovic, J. Pasulka, Z. Zdrahal, R. Stefl, S. Vanacova, RBM7 subunit of the NEXT complex binds U-rich sequences and targets 3’-end extended forms of snRNAs, Nucleic Acids Res, 43 (2015) 4236–4248.

[57] C. Maris, C. Dominguez, F.H. Allain, The RNA recognition motif, a plastic RNA-binding platform to regulate post-transcriptional gene expression, FEBS J, 272 (2005) 2118–2131.

[58] S. Falk, K. Finogenova, M. Melko, C. Benda, S. Lykke-Andersen, T.H. Jensen, E. Conti, Structure of the RBM7-ZCCHC8 core of the NEXT complex reveals connections to splicing factors, Nat Commun, 7 (2016) 13573.

[59] N. Sofos, M.B. Winkler, D.E. Brodersen, RRM domain of human RBM7: purification, crystallization and structure determination, Acta Crystallogr F Struct Biol Commun, 72 (2016) 397–402.

[60] J.F. Watson, J. Garcia-Nafria, In vivo DNA assembly using common laboratory bacteria: A re-emerging tool to simplify molecular cloning, J Biol Chem, 294 (2019) 15271–15281.

[61] S. Raran-Kurussi, S. Cherry, D. Zhang, D.S. Waugh, Removal of Affinity Tags with TEV Protease, Methods Mol Biol, 1586 (2017) 221–230.

[62] S. Duvaud, C. Gabella, F. Lisacek, H. Stockinger, V. Ioannidis, C. Durinx, Expasy, the Swiss Bioinformatics Resource Portal, as designed by its users, Nucleic Acids Res, 49 (2021) W216–W227.

[63] Y. Shimizu, A. Inoue, Y. Tomari, T. Suzuki, T. Yokogawa, K. Nishikawa, T. Ueda, Cell-free translation reconstituted with purified components, Nat Biotechnol, 19 (2001) 751–755.

[64] M.R. Green, J. Sambrook, Isolation of DNA Fragments from Polyacrylamide Gels by the Crush and Soak Method, Cold Spring Harbor Protocols, 2019 (2019) pdb.prot100479.

[65] P.R. Romero, N. Kobayashi, J.R. Wedell, K. Baskaran, T. Iwata, M. Yokochi, D. Maziuk, H. Yao, T. Fujiwara, G. Kurusu, E.L. Ulrich, J.C. Hoch, J.L. Markley, BioMagResBank (BMRB) as a Resource for Structural Biology, Methods Mol Biol, 2112 (2020) 187–218.

[66] Y. Shen, F. Delaglio, G. Cornilescu, A. Bax, TALOS+: a hybrid method for predicting protein backbone torsion angles from NMR chemical shifts, J Biomol NMR, 44 (2009) 213–223.

[67] N.A. Farrow, J.D. Zhang O Fau - Forman-Kay, L.E. Forman-Kay Jd Fau - Kay, L.E. Kay, A heteronuclear correlation experiment for simultaneous determination of 15N longitudinal decay and chemical exchange rates of systems in slow equilibrium.

[68] E.E. Metcalfe, J. Zamoon, D.D. Thomas, G. Veglia, 1H/15N Heteronuclear NMR Spectroscopy Shows Four Dynamic Domains for Phospholamban Reconstituted in Dodecylphosphocholine Micelles, Biophysical Journal, 87 (2004) 1205–1214.

[69] F. Delaglio, S. Grzesiek, G.W. Vuister, G. Zhu, J. Pfeifer, A. Bax, NMRPipe: a multidimensional spectral processing system based on UNIX pipes, J Biomol NMR, 6 (1995) 277–293.

[70] W. Lee, M. Tonelli, J.L. Markley, NMRFAM-SPARKY: enhanced software for biomolecular NMR spectroscopy, Bioinformatics, 31 (2015) 1325–1327.

[71] M.W. Maciejewski, A.D. Schuyler, M.R. Gryk, Moraru, II, P.R. Romero, E.L. Ulrich, H.R. Eghbalnia, M. Livny, F. Delaglio, J.C. Hoch, NMRbox: A Resource for Biomolecular NMR Computation, Biophys J, 112 (2017) 1529–1534.

[72] A. Busch, M. Bruggemann, S. Ebersberger, K. Zarnack, iCLIP data analysis: A complete pipeline from sequencing reads to RBP binding sites, Methods, 178 (2020) 49–62.

[73] I. Huppertz, J. Attig, A. D’Ambrogio, L.E. Easton, C.R. Sibley, Y. Sugimoto, M. Tajnik, J. Konig, J. Ule, iCLIP: protein-RNA interactions at nucleotide resolution, Methods, 65 (2014) 274–287.

[74] http://www.bioinformatics.babraham.ac.uk/projects/fastqc/.

[75] https://www.bioinformatics.babraham.ac.uk/projects/trim_galore/.

[76] M. Briese, N. Haberman, C.R. Sibley, R. Faraway, A.S. Elser, A.M. Chakrabarti, Z. Wang, J. Konig, D. Perera, V.O. Wickramasinghe, A.R. Venkitaraman, N.M. Luscombe, L. Saieva, L. Pellizzoni, C.W.J. Smith, T. Curk, J. Ule, A systems view of spliceosomal assembly and branchpoints with iCLIP, Nat Struct Mol Biol, 26 (2019) 930–940.

[77] https://github.com/alexdobin/STAR.

[78] A.R. Quinlan, I.M. Hall, BEDTools: a flexible suite of utilities for comparing genomic features, Bioinformatics, 26 (2010) 841–842.

[79] K. Darty, A. Denise, Y. Ponty, VARNA: Interactive drawing and editing of the RNA secondary structure, Bioinformatics, 25 (2009) 1974–1975.

[80] M.R. Puno, C.D. Lima, Structural basis for MTR4-ZCCHC8 interactions that stimulate the MTR4 helicase in the nuclear exosome-targeting complex, Proc Natl Acad Sci U S A, 115 (2018) E5506–E5515.

[81] S.W. Olson, A.W. Turner, J.W. Arney, I. Saleem, C.A. Weidmann, D.M. Margolis, K.M. Weeks, A.M. Mustoe, Discovery of a large-scale, cell-state-responsive allosteric switch in the 7SK RNA using DANCE-MaP, Mol Cell, 82 (2022) 1708–1723 e1710.

[82] L. Luo, L.Y. Chiu, A. Sugarman, P. Gupta, S. Rouskin, B.S. Tolbert, HnRNP A1/A2 Proteins Assemble onto 7SK snRNA via Context Dependent Interactions, J Mol Biol, 433 (2021) 166885.

[83] R. Van Damme, K. Li, M. Zhang, J. Bai, W.H. Lee, J.D. Yesselman, Z. Lu, W.A. Velema, Chemical reversible crosslinking enables measurement of RNA 3D distances and alternative conformations in cells, Nature Communications, 13 (2022) 911.

[84] G.M. Daubner, A. Clery, F.H. Allain, RRM-RNA recognition: NMR or crystallography…and new findings, Curr Opin Struct Biol, 23 (2013) 100–108.

[85] N. Bhattacharjee, P. Biswas, Position-specific propensities of amino acids in the beta-strand, BMC Struct Biol, 10 (2010) 29.

[86] C. Barrandon, F. Bonnet, V.T. Nguyen, V. Labas, O. Bensaude, The transcription-dependent dissociation of P-TEFb-HEXIM1-7SK RNA relies upon formation of hnRNP-7SK RNA complexes, Mol Cell Biol, 27 (2007) 6996–7006.

