## Supplemental File 1 for "C-terminal determinants for RNA binding motif 7 protein stability and RNA recognition"

**Table of contents:**

**Table S1.** Extinction coefficients of RBM7 protein constructs

**Table S2.** Nucleotide sequences of the 7SK SL3 RNA constructs

**Table S3.** Protein and RNA amounts used in sample preparation for EMSA

**Figure S1.** SDS-PAGE gel of HT-RBM7$^{FL}$

**Figure S2.** Stability of HT-RBM7$^{91}$ and HT-RBM7$^{101}$ over time

**Figure S3.** Survey of determined structures of human RBM7 RRM

**Figure S4.** Intermolecular interactions observed in the pentameric unit cell of the X-ray crystal structure PDB ID 5IQQ

**Table S1.** The extinction coefficients of RBM7 protein constructs calculated using Expasy ProtParam website.

| Protein construct | Extinction coefficient ($M^{-1}$ $cm^{-1}$) |
|---|---|
| HT-$RBM7^{FL}$ | 102220 |
| HMT-$RBM7^{91}$ | 70820.0 |
| HT-$RBM7^{86}$ | 4470.00 |
| HT-$RBM7^{91}$ | 4470.00 |
| HT-$RBM7^{101}$ | 5960.00 |
| HGT-$RBM7^{101}$ | 15930.0 |
| $RBM7^{101}$ | 4470.00 |

**Table S2.** The nucleotide sequences of the 7SK SL3 RNA constructs arranged from 5'- 3'. The underlined nucleotides indicate base pairing.

| RNA constructs | |
|---|---|
| $SL3_{210-264}$ | CCCUGCUAGAACCUCCAAACAAGCUCUCAAGGUCCAUUUGUAGGAGAACGUAGGG |
| $SL3_{235-251}$ | CUCAAGGUCCAUUUGUA |
| $SL3_{244-262}$ | CAUUUGUAGGAGAACGUAG |

**Table S1.** The amounts of protein and RNA used in sample preparation for EMSA experiments

| Sample | Molar ratios (RNA:protein) | RNA (pmol) | protein (pmol) |
|---|---|---|---|
| **1** | Protein only | 0 | 1000 |
| **2** | 1:0 | 50 | 0 |
| **3** | 1:2 | 50 | 100 |
| **4** | 1:4 | 50 | 200 |
| **5** | 1:6 | 50 | 300 |
| **6** | 1:8 | 50 | 400 |
| **7** | 1:10 | 50 | 500 |
| **8** | 1:12 | 50 | 600 |
| **9** | 1:15 | 50 | 750 |
| **10** | 1:20 | 50 | 1000 |

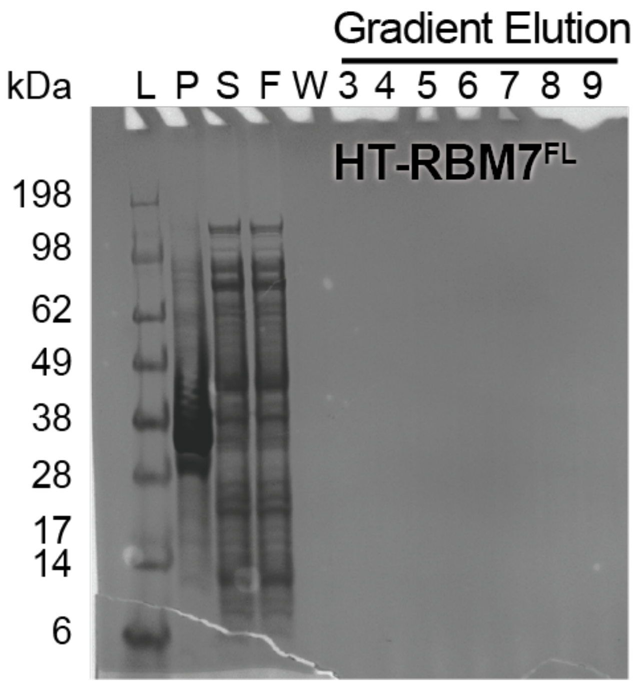


**Figure S1.** SDS-PAGE gel of HT-RBM7$^{FL}$ cell pellet, supernatant of cell lysate, flow-through of the lysate from Ni-NTA column, column wash with buffer R and collected fractions.

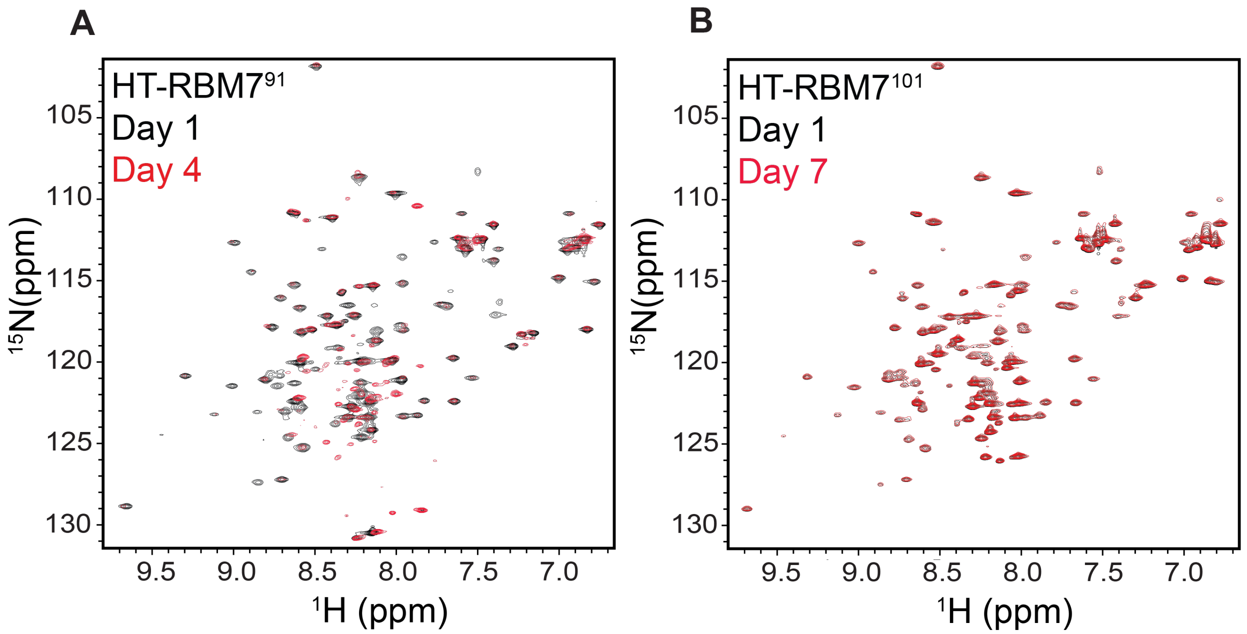


**Figure S2.** $^{1}H$-$^{15}N$ HSQC spectra overlay of A) HT-RBM7$^{91}$ and b) HT-RBM7$^{101}$ over time. All spectra were collected at 600 MHz and 20 ºC in RBM7 storage buffer.

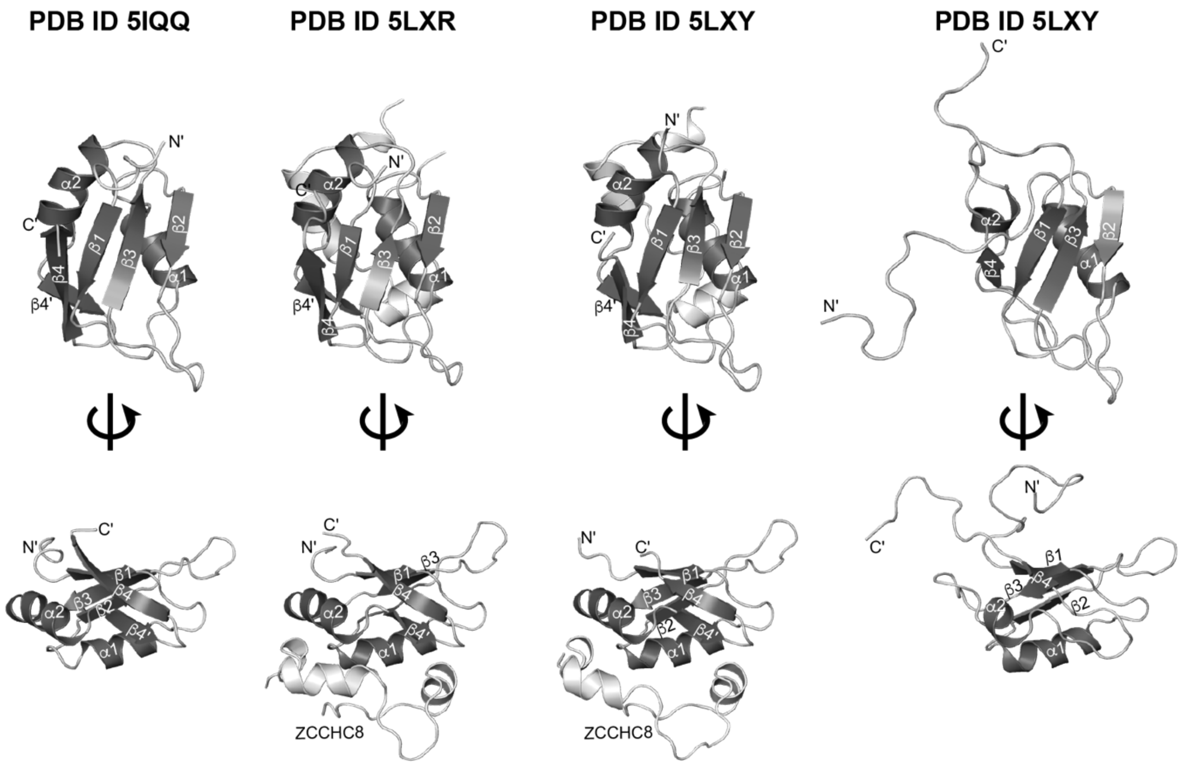


**Figure S3.** Survey of determined structures of human RBM7 RRM.

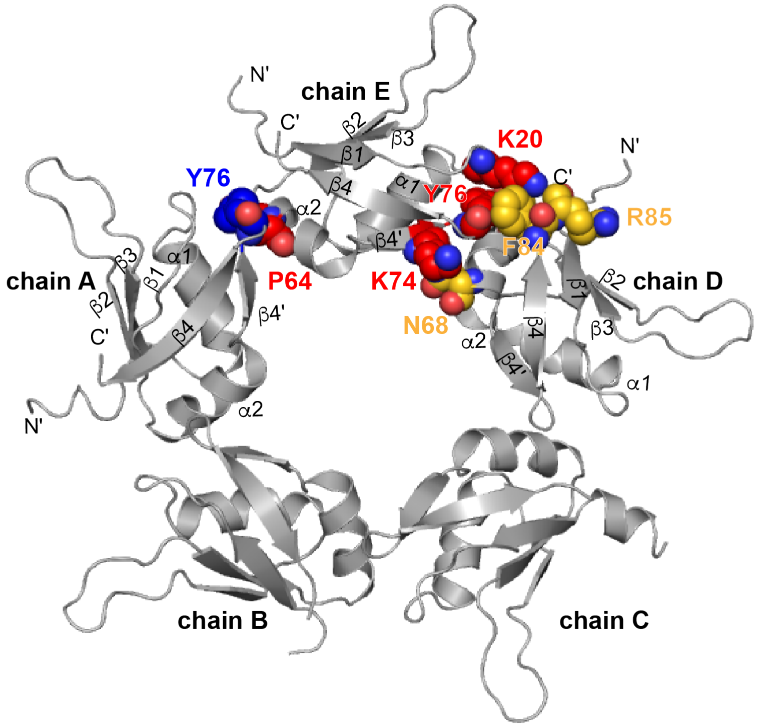


**PDB ID 5IQQ**

**Figure S4.** Intermolecular interactions observed in the pentameric unit cell of the X-ray crystal structure PDB ID 5IQQ. Residues involved in intermolecular crystal contacts in chain E shown in red, chain D shown in gold, and chain A shown in blue.
